# The first *Antechinus* reference genome provides a resource for investigating the genetic basis of semelparity and age-related neuropathologies

**DOI:** 10.1101/2020.09.21.305722

**Authors:** Parice A. Brandies, Simon Tang, Robert S.P. Johnson, Carolyn J. Hogg, Katherine Belov

## Abstract

Antechinus are a genus of mouse-like marsupials that exhibit a rare reproductive strategy known as semelparity and also naturally develop age-related neuropathologies similar to those in humans. We provide the first annotated antechinus reference genome for the brown antechinus (*Antechinus stuartii*). The reference genome is 3.3Gb in size with a scaffold N50 of 73Mb and 93.3% complete mammalian BUSCOs. Using bioinformatic methods we assign scaffolds to chromosomes and identify 0.78Mb of Y-chromosome scaffolds. Comparative genomics revealed interesting expansions in the NMRK2 gene and the protocadherin gamma family, which have previously been associated with aging and age-related dementias respectively. Transcriptome data displayed expression of common Alzheimer’s related genes in the antechinus brain and highlight the potential of utilising the antechinus as a future disease model. The valuable genomic resources provided herein will enable future research to explore the genetic basis of semelparity and age-related processes in the antechinus.

## Context

Antechinus are a genus of small, carnivorous, dasyurid marsupials that are distributed throughout Australia and New Guinea, and exhibit a rare reproductive strategy known as semelparity. Semelparous species reproduce only once in a lifetime [1]. Although this reproductive strategy is common among bacteria, plant and invertebrate species [2], it is rarely seen in mammalian species and is restricted to didelphid and dasyurid marsupials [3, 4]. During the annual breeding season, male antechinus undergo an extreme shift in resource allocation from survival to reproduction, resulting in a complete die-off of all males in the weeks following mating [1, 5-7]. Increased levels of plasma corticosteroid assist antechinus males in utilising their energy reserves to maximise reproductive potential during the breeding season [4]. However, elevation of these corticosteroids results in total immune system collapse leading to gastrointestinal haemorrhage, parasite/pathogen invasion and death [6, 8]. It is currently unknown how semelparity is controlled at the genetic level in the antechinus.

The antechinus has also been proposed as a model species for the physiology of dementias associated with aging such as Alzheimer’s disease (AD) [3, 9, 10]. Primarily characterised by the formation of amyloid-β plaques and neurofibrillary tangles in the brain, AD is a progressive neurodegenerative disease that is predicted to affect more than 100 million people by 2050 [11]. Traditionally, transgenic mouse models have been utilised to study AD [12-14]; however, mice do not naturally develop β-amyloid plaques and neurofibrillary tangles [15, 16]. Both of these have been found to develop naturally in mature male and female antechinus, particularly after the breeding season [9, 10].

Antechinus also possess a number of characteristics that could make them an ideal model organism including: a small body size, short lifespan, production of large numbers of offspring and the ability to be easily maintained in captivity [6, 17, 18]. Creating a reference genome for the antechinus and understanding whether there is expression of key AD-related genes in the antechinus’ brain is a key first step in determining their suitability as a future disease model for AD in humans.

Here we present an annotated reference genome for the brown antechinus (*Antechinus stuartii*). We use a bioinformatic approach [19] to provide a more complete characterisation of the Y chromosome which is currently poorly annotated in marsupials, due to its heterochromatic, highly repetitive nature and small size [20]. We also call and annotate phased genome-wide SNVs (single nucleotide variants) and structural variants, and use comparative genomics to identify rapidly evolving gene families. Finally, we characterise variation in a variety of genes that have previously been associated with AD and evaluate the expression of these genes in the antechinus transcriptome.

The annotated genome and other genomic resources provided herein provide a powerful foundation for studying semelparity and neurodegeneration as well as showcasing the potential hidden within the genomes of Australia’s unique biodiversity.

## Methods

### Sample Collection

Using a standard Elliot trapping procedure (University of Sydney Animal Ethics: 2018/1438) [21], one male and one female adult brown antechinus were trapped in June 2019 at Lane Cove National Park, NSW. Individuals were euthanased using pentobarbitone (60mg/mL) and samples were collected immediately after death. Blood samples were collected in RNAprotect® Animal Blood Tubes and stored at 4°C. Tissue samples were either flash frozen in liquid nitrogen (genomic DNA extraction) or placed in RNAlater (transcriptomic RNA extraction) and stored at 4°C overnight before long-term storage at - 80°C.

### Genome Assembly

DNA was extracted from female and male skeletal muscle tissue using the Circulomics Nanobind HMW DNA kit and quantified using a Qubit dsDNA BR (Broad Range) assay and pulse field gel electrophoresis. 10X Genomics linked-read sequencing libraries were prepared at the Ramaciotti Centre for Genomics (Sydney, NSW, Australia) and sequenced on a NovaSeq 6000 S1 flowcell using 150bp PE reads. *De novo* genome assembly was performed for both sexes independently with Supernova v2.1.1 (RRID:SCR_016756) [22] using all reads, obtaining approximately 75x raw coverage and 55x effective (deduplicated) coverage. BBTools v38.73 (RRID:SCR_016968) [23] was used to generate assembly statistics and BUSCO (RRID:SCR_015008) [24] analysis was performed with both v3.0.2 (4,104 mammalian BUSCOs) and v 4.0.6 (9,226 mammalian BUSCOs).

### Chromosome Assignment and Y Chromosome Analysis

Putative chromosome assignment of the male assembly was achieved by mapping the male scaffolds to the chromosome-length reference genome of the closely-related Tasmanian devil (*Sarcophilus harrisii*) available on NCBI (RefSeq assembly mSarHar1.11, RRID:SCR_003496) [25] using nucmer v4.0.0beta2 (RRID:SCR_018171) [26] and filtering the output using custom bash scripts. Due to the lack of complete Y chromosome sequence in the Tasmanian devil reference genome, additional Y chromosome scaffolds were identified using an AD-ratio (average depth ratio) approach [19] and confirmed through BLAST searches of known marsupial Y genes.

Firstly, both the male and female 10x reads were trimmed to remove the 10x Chromium barcode and low-quality sequence using FastQC v0.11.5 (RRID:SCR_014583) [27] and BBTools (RRID:SCR_016968). Male and female trimmed reads were aligned to the male genome assembly separately using BWA (Burrows-Wheeler Aligner) v0.7.17-r1188 (RRID:SCR_010910) [28], duplicates were removed using samblaster v0.1.24 (RRID:SCR_000468) [29] and alignments with quality scores <20 were removed with samtools v1.10 (RRID:SCR_002105) [30]. The output file was converted to bam format, sorted and indexed with samtools and average coverage statistics were generated using Mosdepth v0.2.6 (RRID:SCR_018929) [31]. The AD-ratio of each scaffold was calculated for each scaffold whereby a normalized ratio of female reads to male reads should result in a value of ∼1 for autosomal scaffolds (as both the male and female should have similar levels of coverage at these regions), a value of ∼2 for X chromosome scaffolds (as females should have double the coverage at these regions due to them possessing two X chromosomes) and a value of ∼0 for Y chromosomes (as females should have no coverage at these regions due to the lack of a Y chromosome) [19].

In order to improve our confidence in the scaffolds assigned as putatively male using the AD-ratio approach, we used BLAST v2.6.0 (RRID:SCR_004870) [32, 33] to map 20 known marsupial Y genes and their autosomal or X homologs (if available) from a previous study [34]) against the male antechinus assembly. Scaffolds with an AD-ratio <0.3 and strong BLAST matches (1e^-10^) to marsupial Y genes (but not the respective X chromosome homologs), were deemed as belonging to the Y chromosome.

### Transcriptome Assembly, Annotation and Analysis

Total RNA (excluding miRNA) was extracted from blood using the Qiagen RNeasy Protect Animal Blood Kit, and from tissues using the Qiagen RNeasy Mini Kit with quantification performed using the Agilent Bioanalyzer RNA 6000 Nano Kit. TruSeq Stranded mRNA-seq library preparation was performed on male and female spleen, brain, adrenal gland and reproductive tissues (ovary/testis) at the Ramaciotti Centre for Genomics (Sydney, NSW, Australia), and sequenced as 150bp PE reads on a NovaSeq 6000 SP flowcell. RNA-seq reads were quality trimmed and assembled *de novo* to create a global transcriptome assembly using Trinity v2.10.0 (RRID:SCR_013048) [35, 36] with default Trimmomatic (RRID:SCR_011848) [37] and Trinity parameters. Trinity’s TrinityStats.pl script was used for general assembly statistics, representation of full-length reconstructed protein-coding genes was examined by Swiss-Prot (RRID:SCR_002380) [38] BLAST searches (RRID:SCR_004870), and completeness was assessed using BUSCO (RRID:SCR_015008) v3 and v4. Trimmed reads were mapped back to the assembly using bowtie2 v2.3.5.1 (RRID:SCR_005476) [39] to determine read representation. Transcript abundance for each tissue type was estimated using Trinity (RRID:SCR_013048) and Salmon v1.0.0 (RRID:SCR_017036) [40] to create a cross-sample TMM normalised matrix of expression values [41, 42]. Finally, the ExN50 statistic was calculated using the normalised expression data. This statistic calculates the N50 for the most highly expressed genes thereby excluding any lowly expressed contigs which are often very short (due to low read coverage preventing assembly of complete transcripts) and hence provides a more useful indicator of transcriptome quality than the standard N50 metric [36].

Functional annotation of the global transcriptome was performed using Trinotate v3.2.0 (RRID:SCR_018930) [43]. Briefly, TransDECODER v5.5.0 (RRID:SCR_017647) was used to identify candidate coding regions within the Trinity transcripts. Blast searches of the TransDECODER peptides and Trinity transcripts were performed against the Swiss-Prot (RRID:SCR_002380) database and the Tasmanian devil reference genome annotations from NCBI (RefSeq assembly mSarHar1.11, RRID:SCR_003496) [25] with an e-value cut-off of 1e^-5^. HMMER v3.2.0 (RRID:SCR_005305) [44] was used to identify conserved protein domains with the Pfam (RRID:SCR_004726) [45] database, SignalP v4.1 (RRID:SCR_015644) [46] was used to predict signal peptides and RNAmmer v1.2 (RRID:SCR_017075) [47] was used to detect any ribosomal RNA contamination. The results from the above were loaded into a SQLite3 (RRID:SCR_017672) database with gene ontology (GO) (RRID:SCR_002811) and KEGG (RRID:SCR_012773) terms assigned based on the Swiss-Prot (RRID:SCR_002380) annotations.

### Repeat Identification and Genome Annotation

A custom repeat database was generated with RepeatModeler v2.0.1 (RRID:SCR_015027) [48] and repeats (excluding low complexity regions and simple repeats) were masked with RepeatMasker (RRID:SCR_012954) v4.0.6 [49]. Genome annotation was performed using Fgenesh++ v7.2.2 (RRID:SCR_018928) [50-52] using optimised gene finding parameters of the closely related Tasmanian devil (*Sarcophilus harrisii*) with mammalian general pipeline parameters. Transcripts representing the longest protein for each trinity “gene” were extracted from the trinity and trinotate output files for mRNA-based predictions with a custom bash script using seqtk v1.3 (RRID:SCR_018927) and seqkit v0.10.1 (RRID:SCR_018926) [53]. A high-quality non-redundant metazoan protein dataset from NCBI was used for homology-based predictions using the “prot_map” method. *Ab initio* predictions were performed in regions where no genes were predicted by other methods (i.e. mRNA mapping or protein homology). The predicted protein-coding sequences were used in BLAST (RRID:SCR_004870) searches against the Swiss-Prot (RRID:SCR_002380) database with an e-value cut-off of 1e^-5^ to identify genes with matches to known high quality proteins from other species.

### Variant Annotation

The male reference genome was altered following the 10x Genomics Long Ranger (RRID:SCR_018925) [54] software recommendations of a maximum 500 fasta sequences as follows: scaffolds <50kb were extracted and concatenated with gaps of 500 N’s and then added to the main genome fasta file as a single scaffold and scaffolds ≥50kb (428 scaffolds) were listed in the primary_contigs.txt file. A BED file of the assembly gaps was created using faToTwoBit and twoBitinfo (RRID:SCR_005780) [55] to generate the sv_blacklist.bed file. Male and female 10x reads were aligned to the altered male 10x reference genome with whole-genome SNVs, indels and structural variants called and phased using Long Ranger v2.2.2 (RRID:SCR_018925) [54] with default parameters. Male and female VCF files were merged with bcftools v1.10.1 (RRID:SCR_002105) [30] and variants were annotated using ANNOVAR v20180416 (RRID:SCR_012821) [56, 57].

### Gene Family Analysis

Gene ontology (GO) annotation (using the generic GO slim subset) was performed on antechinus proteins based on Swiss-Prot matches using GOnet [58] (RRID:SCR_018977) to identify genes associated with key biological functions.

To identify any rapidly evolving gene families in the antechinus, proteomes from six other target species (Tasmanian devil, koala, opossum, human, mouse and platypus) were downloaded from NCBI (RRID:SCR_003496) [25] and the longest isoform for each gene was extracted using custom scripts. Protein sequences from the antechinus Fgenesh++ annotation were also extracted and OrthoFinder v2.4.0 (RRID:SCR_017118) [59, 60] was used to identify orthogroups between the 7 target species. CAFE v5 (RRID:SCR_018924) [61, 62] was run on the output data from OrthoFinder (RRID:SCR_017118) using an error model to account for genome assembly error and estimating multiple lambda’s (gene family evolution rates) for monotremes, marsupials and eutherians. Significant expansions and contractions within the antechinus branch were examined to identify any interesting patterns.

### Alzheimer’s Genes Analysis

Literature searches using the search terms “Alzheimer’s” and “gene”, and mining the human gene database GeneCards [63] using the keyword “Alzheimer’s” were used to identify forty of the most common genes that have previously been associated with Alzheimer’s disease in humans or mice disease models. Human coding sequences (CDS) for the genes of interest were downloaded from Swiss-Prot (RRID:SCR_002380) and were used in BLAST (RRID:SCR_004870) searches against the Fgenesh++ genome annotations to identify the predicted gene sequences within the male antechinus reference genome. The predicted protein sequences were matched against the predicted coding sequences of the global transcriptome using BLAST (RRID:SCR_004870) to identify candidate transcripts and expression of the candidate genes across the sequenced tissues was explored using the TMM-normalised expression matrix. All sequences were aligned to human protein sequences with MUSCLE v3.8.425 [64] in order to determine sequence similarity and identity. SNVs associated with the target genes were explored using the ANNOVAR (RRID:SCR_012821) output.

## Findings

### Genome Assembly

The male and female antechinus genome assemblies were both 3.3Gb in size. Genome contiguity was slightly higher for the male antechinus with a scaffold N50 of 72.7Mb in comparison with the female scaffold N50 of 58.2Mb (Table 1). Both male and female genome assemblies showed completeness scores comparable to the two best marsupial reference genomes currently available (the koala: RefSeq phaCin_unsw_v4.1, and the Tasmanian devil: RefSeq mSarHar1.11), with >90% of the 4,104 version 3 mammalian BUSCO’s and >80% of the 9,226 version 4 mammalian BUSCO’s being complete (Table 1, Supplementary Table 1). The male assembly was chosen to be the reference genome as it showed the highest contiguity and also includes the Y chromosome.

**Table 1.** Comparison of antechinus genome assembly statistics in comparison with the two current highest-quality marsupial genomes.

| SPECIES | ASSEMBLY | GENOME<br>SIZE (Gb) | NO.<br>SCAFFOLDS<br>↓ | NO.<br>CONTIGS<br>↓ | SCAFFOLD<br>N50 (Mb)<br>↑ | CONTIG<br>N50 (Mb)<br>↑ | % GENOME IN<br>SCAFFOLDS ><br>50KB ↑ | COMPLETE<br>MAMMALIAN<br>BUSCO'S V3 (%)<br>↑ | COMPLETE<br>MAMMALIAN<br>BUSCO'S V4 (%)<br>↑ |
| --- | --- | --- | --- | --- | --- | --- | --- | --- | --- |
| **ANTECHINUS**<br>**(M)** | antechinusM_pseudohap2.1 (USYD_AStu_M<sup>1</sup>) | 3.3 | 30876 | 106199 | 72.7 | 0.08 | 96.35 | 93.3 | 81.3 |
| **ANTECHINUS**<br>**(F)** | antechinusF_pseudohap2.1 | 3.3 | 31296 | 107658 | 58.2 | 0.08 | 96.61 | 92.9 | 81.6 |
| **KOALA** | phaCin_unsw_v4.1<sup>1</sup> | 3.2 | - | 1909 | - | 11.59 | 99.11 | 92.3 | 81.6 |
| **TASMANIAN**<br>**DEVIL** | mSarHar1.11<sub>1</sub> | 3.1 | 106 | 445 | 611.3 | 62.34 | 99.97 | 93.8 | 80.9 |

Arrows indicate whether higher or lower numbers are considered better quality.
1. NCBI Assembly ID

### Chromosome Assignment and Y Chromosome Analysis

The *Dasyuridae* family display a high level of karyotypic conservation with all species having almost identical 2n=14 karyotypes [65]. Antechinus chromosomes were therefore bioinformatically assigned by alignment of the male antechinus scaffolds to the chromosome-length Tasmanian devil reference assembly (RefSeq mSarHar1.11). This resulted in 94.3% of the genome being assigned to chromosomes with the remaining 5.7% of the genome being unassigned either due to no matches to the Tasmanian devil genome or due to multiple alignments where there was no best match to a single chromosome (Figure 1a, Supplementary Table 2). The length of assigned antechinus chromosomes was similar to that of the Tasmanian devil as expected (Figure 1b).

**Table 2.** Summary of repeat classes identified and masked in the antechinus reference genome.

| REPEAT CLASS | COUNT | MASKED (BP) | MASKED (%) |
| --- | --- | --- | --- |
| **DNA** |  |  |  |
| CMC-ENSPM | 267774 | 30028201 | 0.91% |
| GINGER-1 | 13763 | 1594788 | 0.05% |
| PIF-HARBINGER | 763 | 204495 | 0.01% |
| TCMAR-TC1 | 7165 | 1616661 | 0.05% |
| TCMAR-TC2 | 3098 | 1745523 | 0.05% |
| TCMAR-TIGGER | 22186 | 4059186 | 0.12% |
| HAT | 744 | 142335 | 0.00% |
| HAT-AC | 2400 | 291924 | 0.01% |
| HAT-CHARLIE | 143304 | 24400026 | 0.74% |
| HAT-TIP100 | 36557 | 6236166 | 0.19% |
| LINE | 6840 | 2038840 | 0.06% |
| CR1 | 301533 | 59092138 | 1.79% |
| DONG-R4 | 12719 | 4935572 | 0.15% |
| L1 | 1117136 | 608623645 | 18.40% |
| L2 | 770053 | 168785105 | 5.10% |
| RTE-BOVB | 98681 | 30352289 | 0.92% |
| RTE-RTE | 64120 | 17729186 | 0.54% |
| **LTR** |  |  |  |
| ERV1 | 19808 | 9033177 | 0.27% |
| ERVK | 56462 | 49884792 | 1.51% |
| ERVL | 2556 | 1297101 | 0.04% |
| GYPSY | 4842 | 1375235 | 0.04% |
| **SINE** |  |  |  |
| 5S-DEU-L2 | 4816 | 270426 | 0.01% |
| ALU | 6938 | 1367052 | 0.04% |
| MIR | 1445092 | 212663300 | 6.43% |
| **OTHER** |  |  |  |
| UNKNOWN | 1070813 | 233112108 | 7.05% |
| SATELLITE | 52562 | 11605904 | 0.35% |
| SNRNA | 382 | 28484 | 0.00% |
| **TOTAL** | 5533107 | 1482513659 | 44.82% |

**Figure 1.**
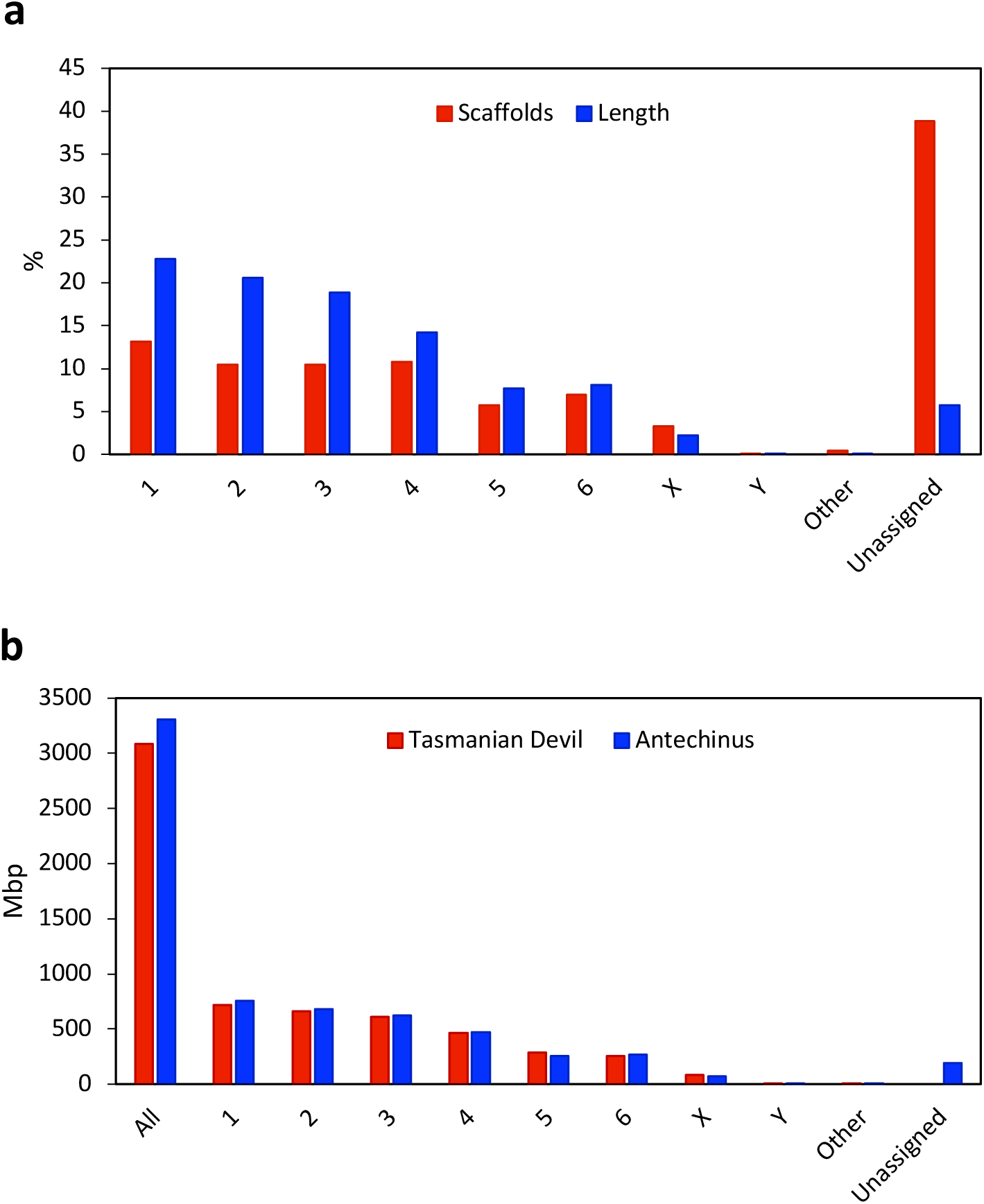
Assignment of antechinus scaffolds to chromosomes by alignment to the Tasmanian devil reference genome. **a)** Proportion (%) of scaffolds (blue) and genome length (red) assigned to chromosomes. **b)** Comparison of length of sequence assigned to each chromosome from the Tasmanian devil reference genome (blue) and the antechinus genome (red). Other represents scaffolds assigned to “unplaced” Tasmanian devil scaffolds and Unassigned represents scaffolds unable to be assigned due to no matches to the Tasmanian devil genome or due to multiple matches where a best hit to a single chromosome was not identified.

The current Tasmanian devil reference genome (RefSeq mSarHar1.11) contains limited Y-chromosome sequence (∼130kb) and so only one antechinus scaffold (scaffold 161317, ∼73kb) was assigned as Y chromosome. To identify further putative Y chromosome scaffolds, we implemented an AD-ratio approach (see [19]). Using this approach 3.1Gb (∼95%) of the male genome was assigned as autosomal, 87Mb (∼2.6%) of the male genome was assigned as X chromosomal and 11.4Mb (0.3%) of the genome was assigned as Y chromosomal (Figure 2). The results from this approach showed that ∼92% of the genome was in agreeance with the chromosome assignment results from mapping the antechinus genome to Tasmanian devil genome with the remaining 8% mainly due to unassigned chromosomes from either method rather than chromosome discrepancies between the two methods (only 0.2% of genome) (Supplementary Table 2).

**Figure 2.**
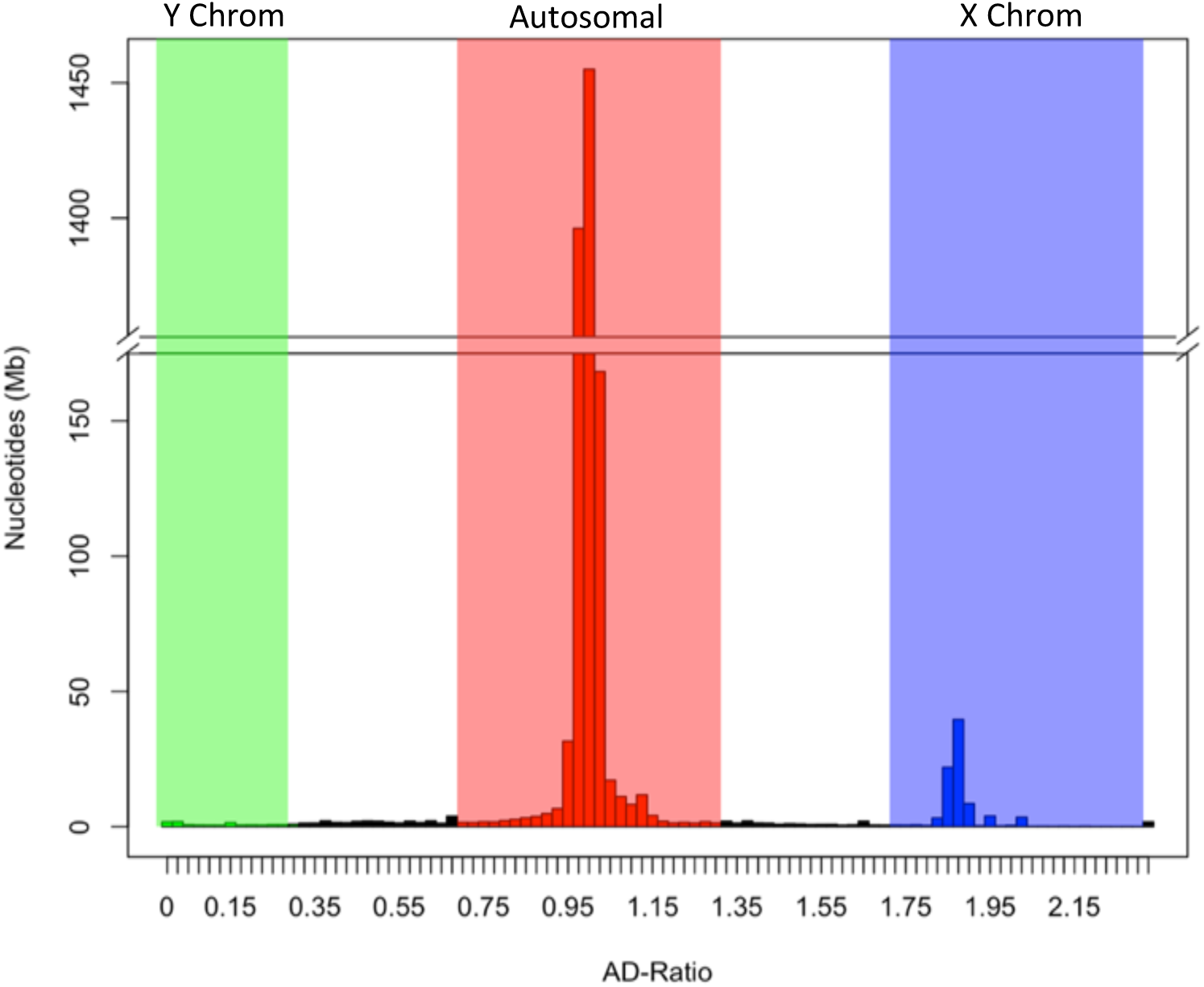
AD-Ratio histogram of antechinus scaffolds. Figure shows the total length of sequence within each 0.025 AD-ratio bin. Scaffolds clustering around an AD-ratio of 0 represent Y-linked sequence (Green), scaffolds clustering around an AD-ratio of 1 represent Autosomal sequence (Red), scaffolds clustering around an AD-ratio of 2 represent X-linked sequence (Blue) and scaffolds between these regions represent unassigned sequence (Black).

In order to identify some high-confidence Y chromosome scaffolds from the putative Y chromosome scaffolds identified with the AD-ratio approach, we aimed to identify scaffolds containing known Y genes and Y-specific transcripts. Out of 20 known marsupial Y chromosome genes from a previous study [34], 13 showed hits to scaffolds with AD-ratios ≤0.01 indicating a high chance they are putative Y chromosome scaffolds. Furthermore, their autosomal, or X chromosome, homologs mapped to different scaffolds providing additional confidence that the scaffolds identified likely contain the Y homolog. Seven of these Y genes were found to be on scaffold 163451, four were located on scaffold 162475 and one was matched to scaffold 161317 (Figure 3). These scaffolds were deemed Y-chromosome scaffolds and comprise 0.78Mb of the genome. They represent the largest amount of Y-chromosome sequence characterized in any marsupial species. The remaining gene (ATRY) displayed multiple partial alignment hits to a number of different antechinus scaffolds and could not be reliably annotated to a single scaffold. A number of other genes were also annotated to these scaffolds by Fgenesh++ annotation including an XK-related protein on scaffold 161317, an AMMECR1-like gene on scaffold 163451 and a HMGB3-like protein on scaffold 162475. Identification and annotation of Y chromosome scaffolds in the antechinus will assist with future research wanting to explore male semelparity and key male-specific reproductive genes.

**Figure 3.**
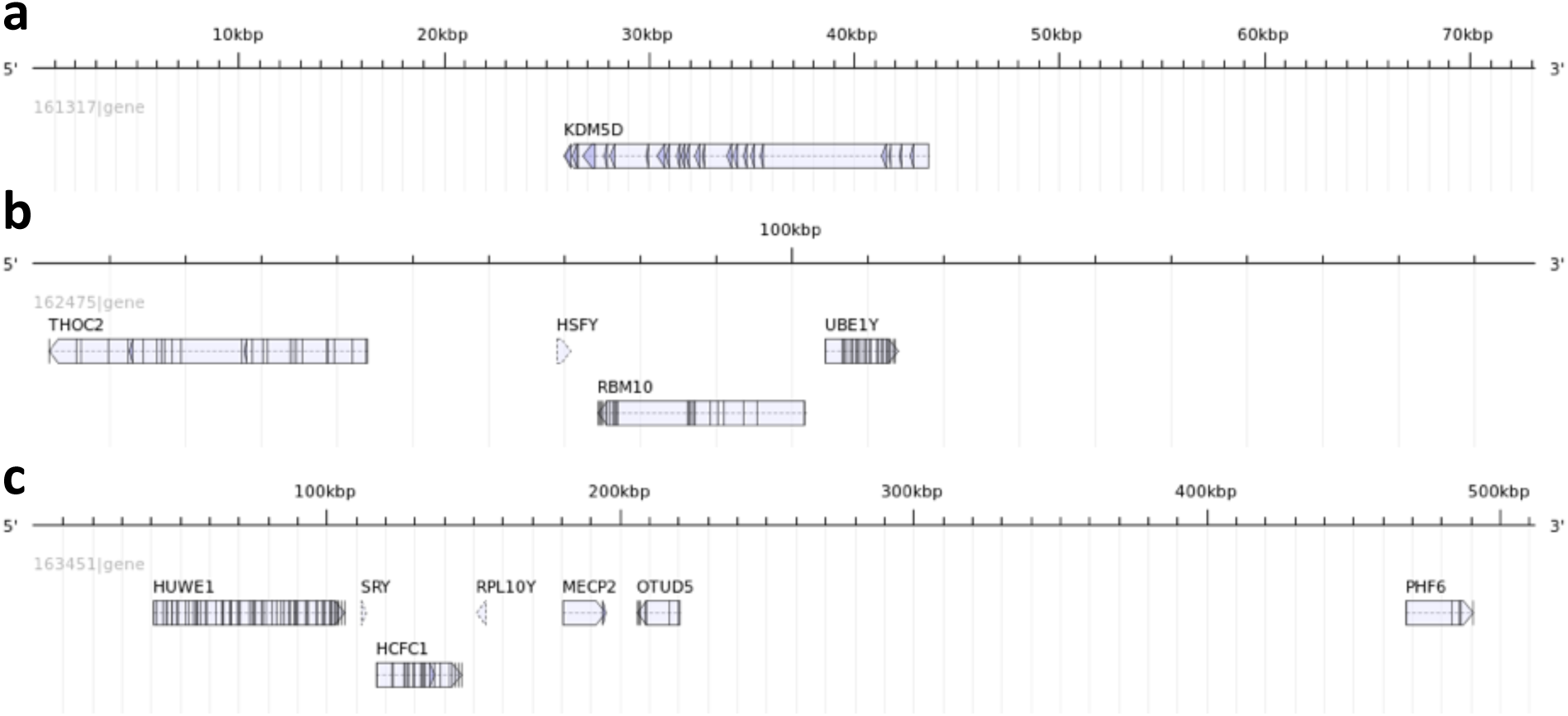
Mapping of known marsupial Y gene homologs on antechinus Y chromosome scaffolds. **a)** Scaffold 161317, **b)** Scaffold 162475, **c)** Scaffold 163451. Figure was created using the AnnotationSketch module from GenomeTools [66].

### Transcriptome Assembly and Annotation

The global antechinus transcriptome assembly of 10 tissues (5 male and 5 female) was composed of 1,296,975 transcripts (1,636,859 including predicted splicing isoforms).

The average contig length was 773bp and the contig N50 was 1,367bp. Considering only the top 95% most highly expressed transcripts gave an ExN50 (a more useful indicator of transcriptome quality) of 3,020bp which is similar to the average mRNA length in humans (3,392bp) [67]. The assembly showed good overall alignment rates of reads from each of the tissues (>96%) with a high percentage mapped as proper pairs (≥89%). The transcriptome assembly exhibited similar completeness to the genome with BUSCO analysis identifying 94% and 84% complete BUSCOs for version 3 and version 4 mammalian datasets respectively (Supplementary Table 1). TransDecoder predicted 296,706 coding regions within the global transcriptome (including predicted splicing isoforms) of which 181,691 (61%) were complete (contained both a start and stop codon) and 159,121 (54%) had BLAST hits to Swiss-Prot. Taking only the longest complete predicted isoform for each gene resulted in 38,829 mRNA transcripts that were used for genome annotation.

### Repeat Identification and Genome Annotation

873 repeat families were identified in the male antechinus genome (Table 2), with 44.82% of the genome being masked as repetitive; a similar repeat content to that of other marsupial and mammalian genomes [68]. A total of 55,827 genes were predicted by Fgenesh++, of which 25,111 had BLAST hits to Swiss-Prot (Supplementary Table 3). This number is similar to that of the 26,856 protein-coding genes annotated in the closely related Tasmanian devil reference genome (RefSeq mSarHar1.11). Of these 25,111 gene annotations, 13,189 were predicted based on transcriptome evidence, 1,286 were predicted based on protein evidence and the remaining were predicted *ab initio* based on trained gene finding parameters.

**Table 3.**
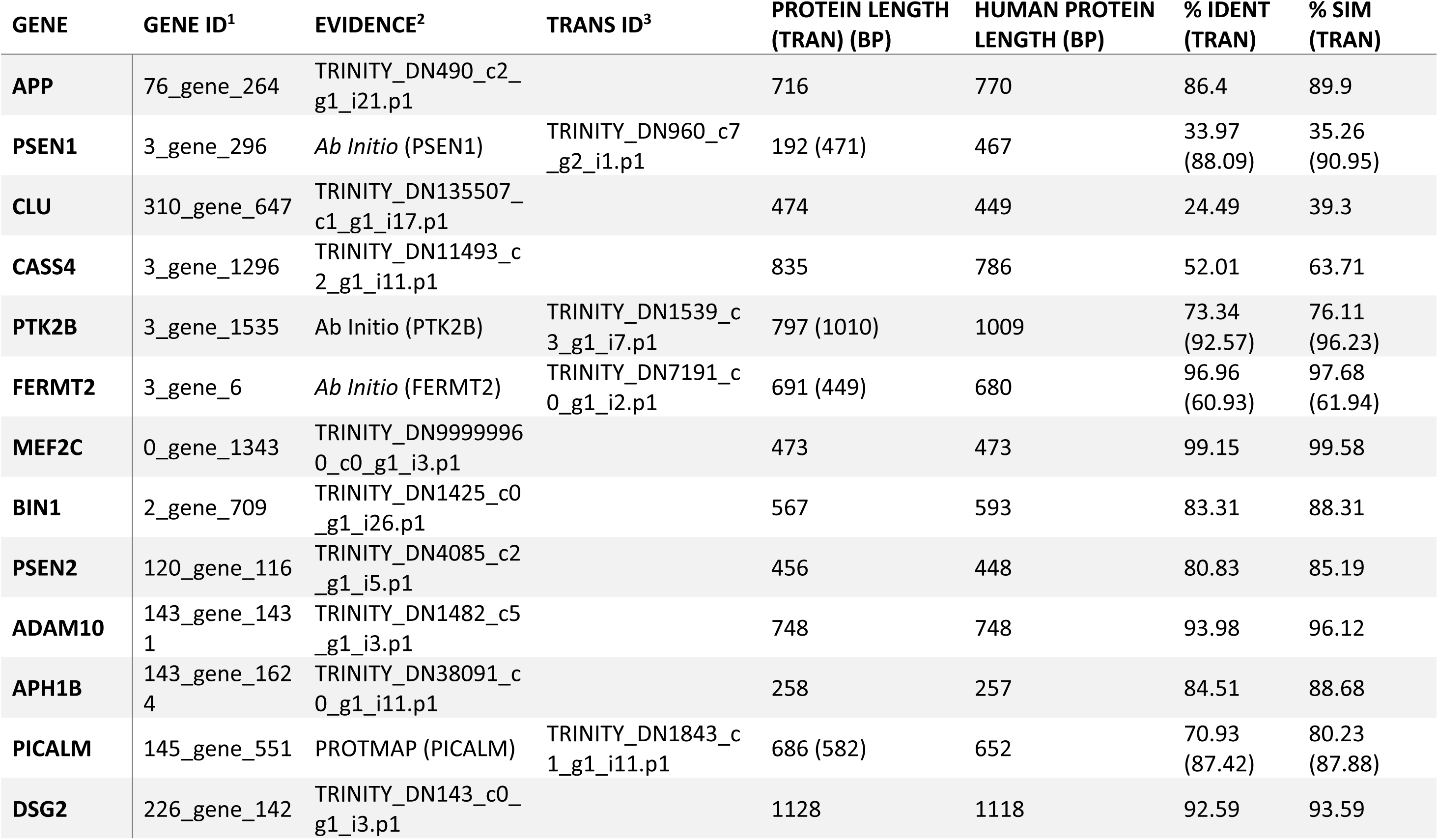
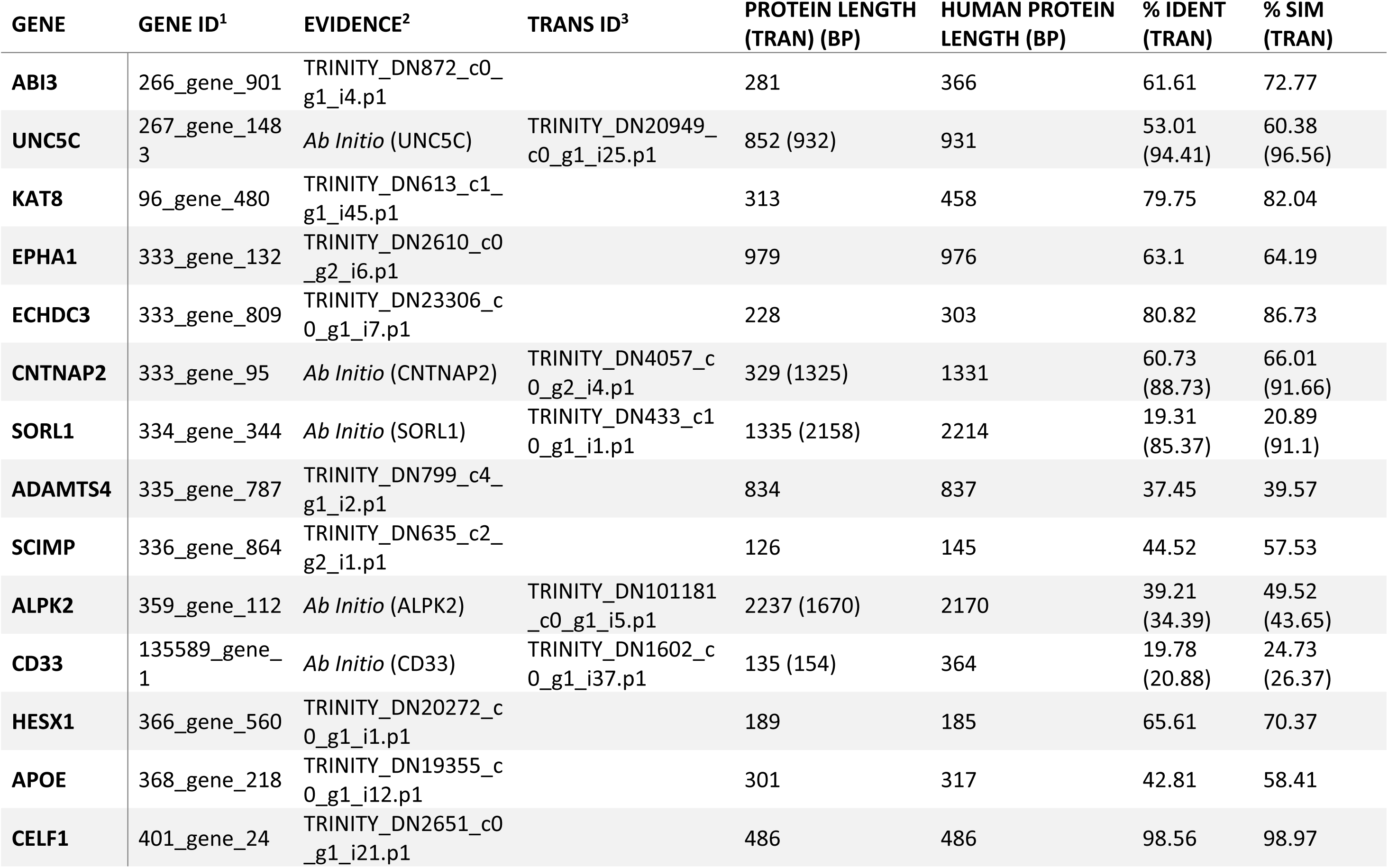
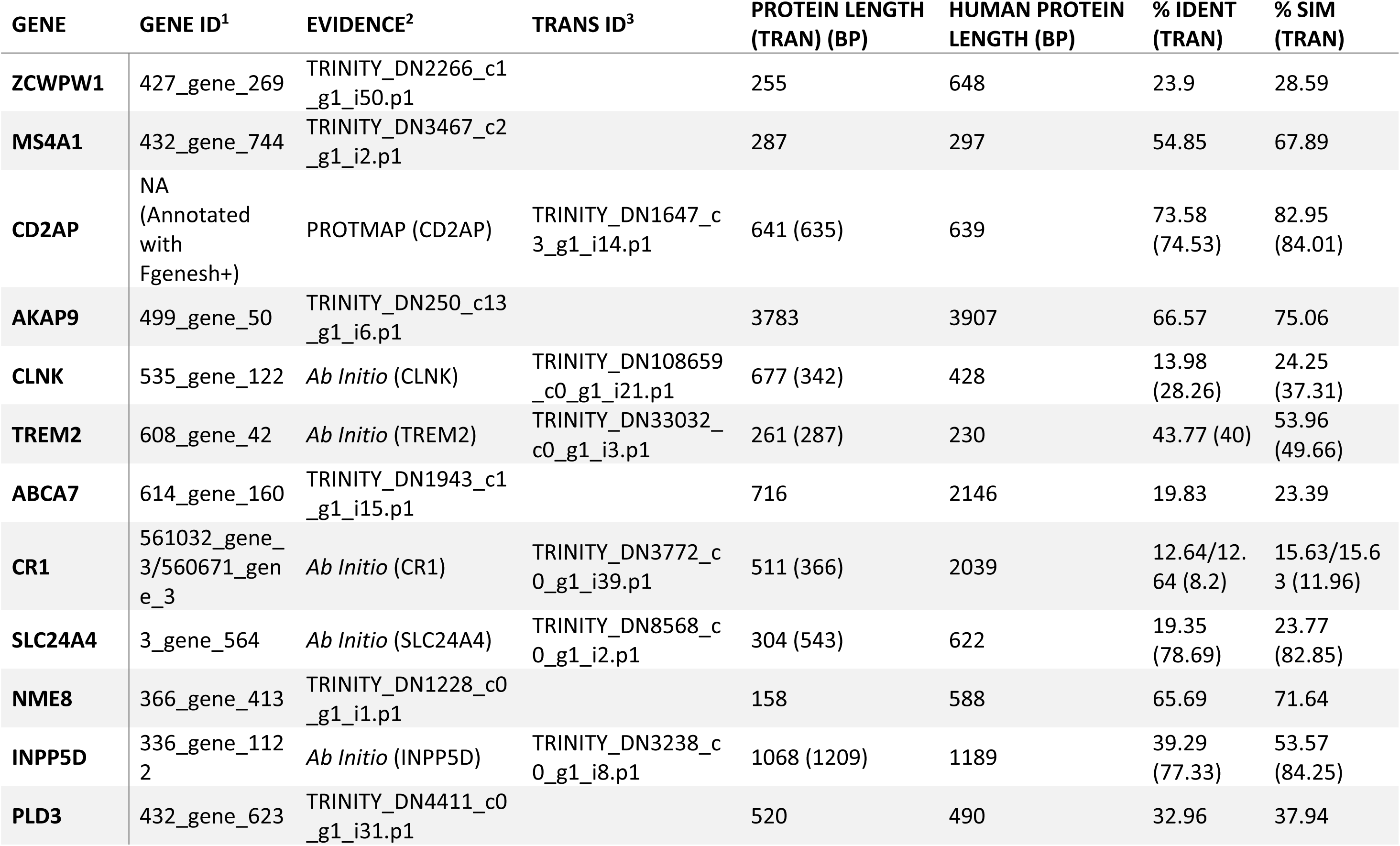
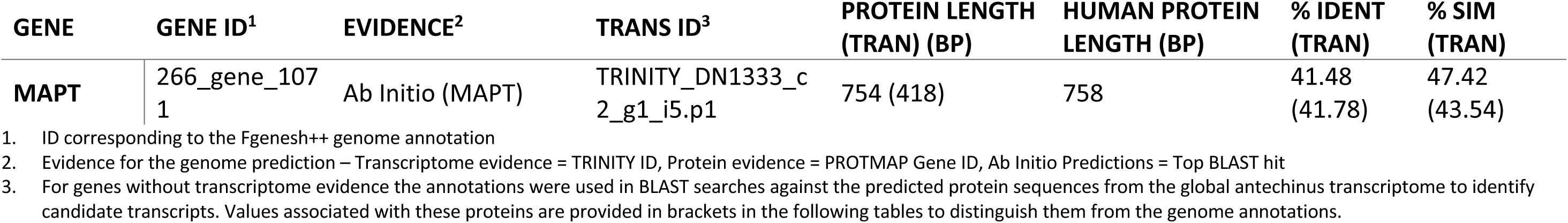
Summary of Alzheimer’s related genes explored in the Antechinus.

### Variant Annotation

The brown antechinus is predicted to be one of the most common and widespread mammalian species in Eastern Australia where it ranges from southern Queensland to southern New South Wales [69, 70]. The large population size and range of *A. stuartii* implies that this species would likely exhibit healthy levels of genomic diversity, though there is currently a lack of genome-wide variation information for any antechinus species. Using the linked-read datasets we identify a total of 9,307,342 SNVs and 2,362,144 indels in the male and 16,291,736 SNVs and 3,818,750 indels in the female; with 5,474,811 SNVs (∼27%) and 1,079,862 indels (∼21%) being genotyped in both individuals. >90% of these variants passed all of the 10X Genomics filters and >99% were phased. Approximately half of the variants were found to be associated with an annotated gene (located within a gene or within 1kb upstream or downstream of a gene) of which 91% were intronic and 2% were exonic (Figure 4a). Within the exonic variants, 58% were nonsynonymous (result in alteration of the protein sequence) and 39% were synonymous (Figure 4b). These results demonstrate considerable genome-wide diversity from just two individuals from the same population. For comparison, just 1,624,852 SNPs (single nucleotide polymorphisms) were identified across 25 individuals of the closely related and endangered Tasmanian devil [71]. Despite the success of *A. stuartii*, other antechinus species, such as the newly-classified and endangered black-tailed dusky antechinus (*A. arktos*), appear in much lower numbers and so may exhibit much lower genome-wide diversity [72]. Most antechinus species diverged in the Pilocene (∼5mya) with the brown antechinus and its close relatives separating more recently in the Pleistocene (∼2.5mya) [73]. Humans and chimpanzees are predicted to have diverged 7-8mya [74] but still share 99% of their DNA [75]. The genetic similarity of human and chimpanzees (which diverged earlier than the antechinus clades) suggests that the annotated antechinus genome and genome-wide variation provided will be a valuable tool to assist with population monitoring and conservation of all species in the antechinus genus.

**Figure 4.**
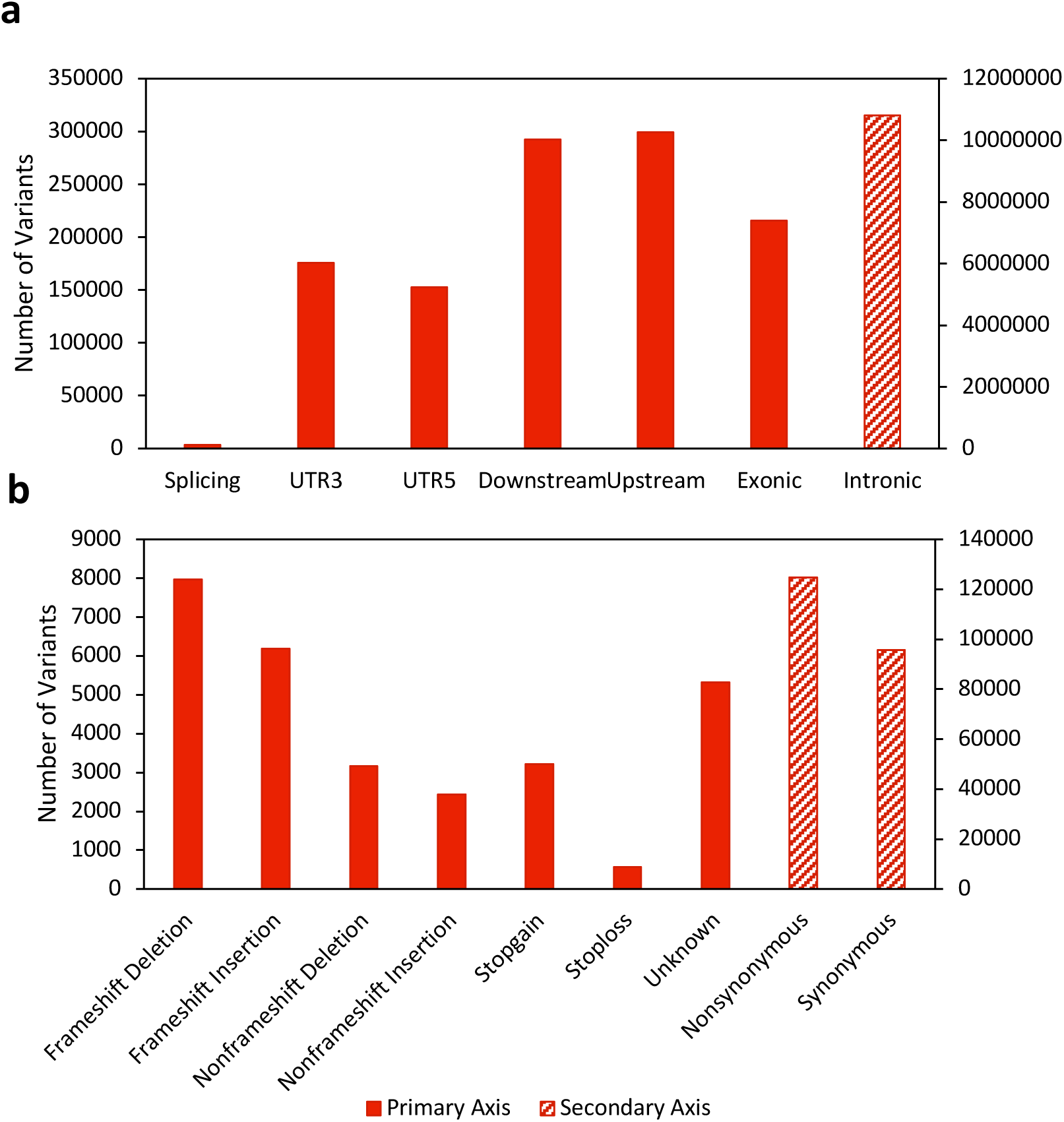
Functional annotation of antechinus variants. **a)** Total number of variants annotated to various gene regions including: Splicing (within a splice site of a gene), UTR3 (3’ untranslated region), UTR5 (5’ untranslated region), Downstream (within 1kb downstream of a gene), Upsteam (within 1kb upstream of a gene), Exonic (within the coding sequence of a gene) and Intronic (within an intron of a gene). **b)** Total number of exonic variants resulting in specific consequences to the protein sequence including: Frameshift Deletion (deletion of one or more nucleotides that results in a frameshift of the coding sequence), Frameshift Insertion (insertion of one or more nucleotides that results in a frameshift of the coding sequence), Nonframeshift Deletion (deletion of one or more nucleotides that does not result in a frameshift of the coding sequence), Nonframeshift Insertion (insertion of one or more nucleotides that does not result in a frameshift of the coding sequence), Stopgain (variation which results in a stop codon being created within the protein sequence), Stoploss (variation which results in a stop codon being lost from the protein sequence), Unknown (variation with an unknown consequence, perhaps due to complex gene structure), Nonsynonymous (a single nucleotide change that does not result in an amino acid change) and Synonymous (a single nucleotide change that results in an amino acid change). Striped bars indicate variant types that are plotted on the secondary Y-axis.

In addition to single nucleotide variants, large structural variants can have a pronounced impact on phenotype and account for a significant amount of the diversity seen between individuals [76, 77]. A few interchromosomal and intrachromosomal rearrangements have been identified in the Dasyuridae family using previous G-banding techniques [78]; however, advancements in sequencing technologies, such as the linked-read approach utilized in the current study, allow for more fine-scale characterisation of structural variants in a cost-effective and reliable manner [79]. Using the linked-read datasets, 700 large, high-quality structural variants were called in the male and 681 were called in the female of which 35% and 25% were copy number variants (CNVs) respectively (Figure 5). Within the intrachromosomal structural variants, 240 in the male, and 191 in the female were found to contain genes, together encompassing 2,401 genes in total (Supplementary Table 4). These findings demonstrate the importance of applying new structural variant identification techniques to explore functional diversity and should be applied more broadly to other Dasyurid species, particularly endangered species such as the Tasmanian devil.

**Figure 5.**
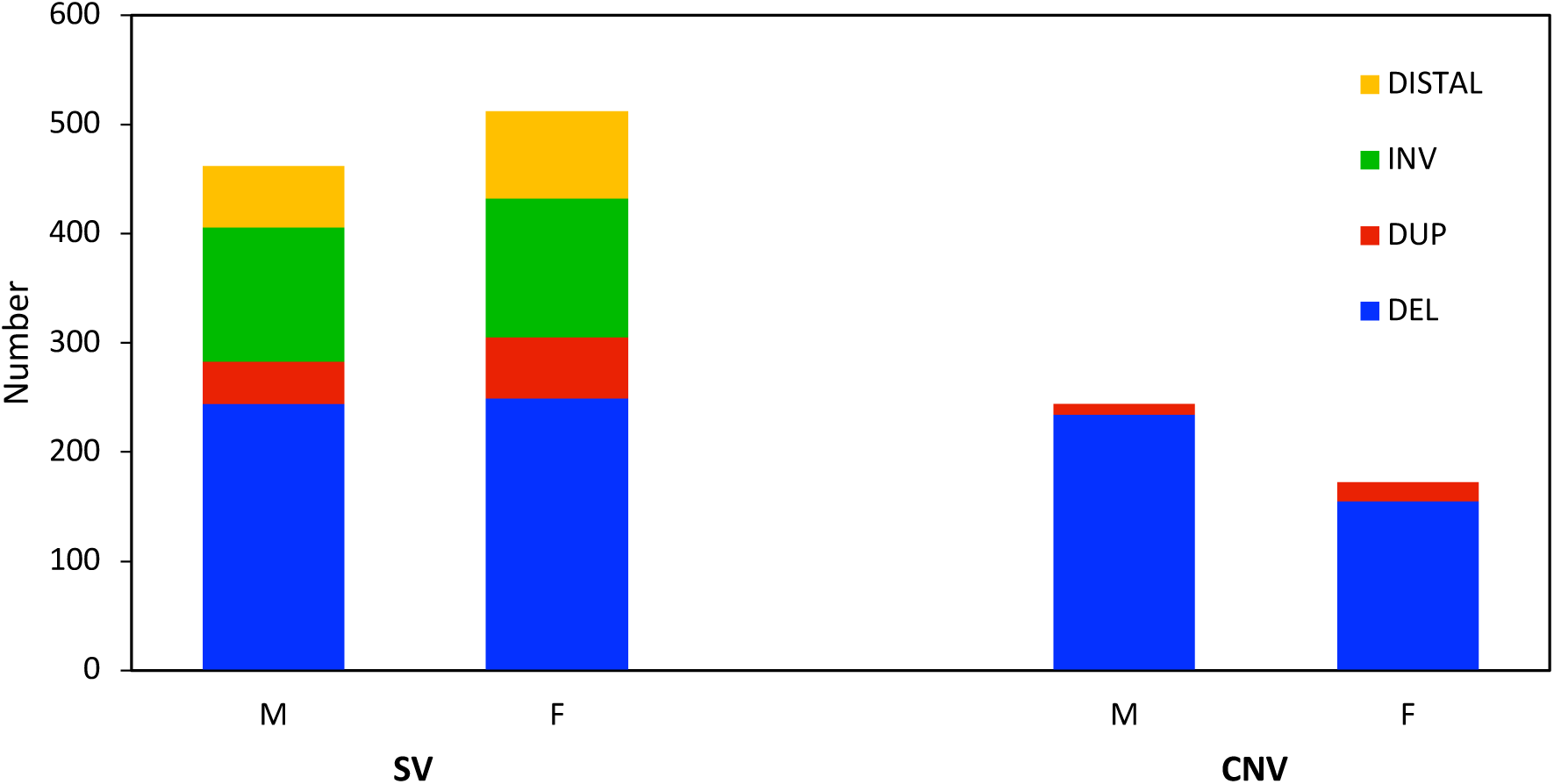
Breakdown of high-quality large structural variants (SVs) and copy number variants (CNVs) in the antechinus. Figure shows both male (M) and female (F) deletions (blue), tandem duplications (red), inversions (green) and distal structural variants (i.e. across two scaffolds, yellow)

### Gene Family Analysis

GO analysis of the antechinus genome annotations based on matches to Swiss-Prot revealed 2,578 of the genes are involved in response to stress, 1,760 are involved in immune system processes and 1,035 are involved in reproduction (Supplementary Table 5). Future studies could use these annotations to design a targeted approach for monitoring the expression of key genes across the breeding season to better understand the interplay between stress, immunity and reproduction in this semelparous species.

To identify any interesting patterns of gene family evolution in the antechinus, proteomes across 7 target species (antechinus, Tasmanian devil, koala, opossum, human, mouse and platypus) were compared and 80.5% of genes were assigned to 19,173 orthogroups of which 12,233 orthogroups had all species present and 9,212 were single-copy orthologs. CAFE identified 282 gene families to be significantly fast evolving (Supplementary Table 6). Of these fast-evolving gene families, a number of significant expansions (<1e-^15^) and contractions were found on the antechinus branch (Supplementary Table 6). Many of these expansions and contractions were found in large, complex gene families including olfactory receptors and immune genes which are notoriously difficult to annotate using automated gene annotation methods, particularly in fragmented assemblies, and so require further investigation and manual curation for confirmation. Two other particularly interesting expansions occurred within the protocadherin gamma (Pcdh-γ) gene family (Orthogroup OG0000022, Supplementary Table 6) and the NRMK2 gene in the antechinus (Orthogroup OG0000350, Supplementary Table 6).

Protocadherins (Pcdhs) belong to the cadherin superfamily and are organised into 3 main gene clusters: α, β and γ [80]. Pcdhs, like all cadherins, are primarily responsible for mediating cell-cell adhesion [81]. Antechinus displayed similar numbers of putative Pcdh-γ genes as humans and mouse (20-21 genes) in comparison to the other marsupials which showed only 6-9 genes in this family, and the platypus only 2 (Figure 6). Pcdh-γ genes specifically have been implicated in neuronal processes [80] and have previously been associated with Alzheimer’s disease [82]. These genes are most highly expressed in the brain in humans and also showed highest levels of expression in the brain and adrenal gland in the antechinus (Supplementary Table 7). It is possible that the expansion of Pcdh-γ genes in the antechinus may be linked to the neuropathological changes that occur in mature antechinus. The α and β Pcdhs were also identified as fast evolving across the 7 target species investigated, with marsupials having lower numbers of genes than eutherians, though there were no large differences in the antechinus branch for these clusters.

**Figure 6.**
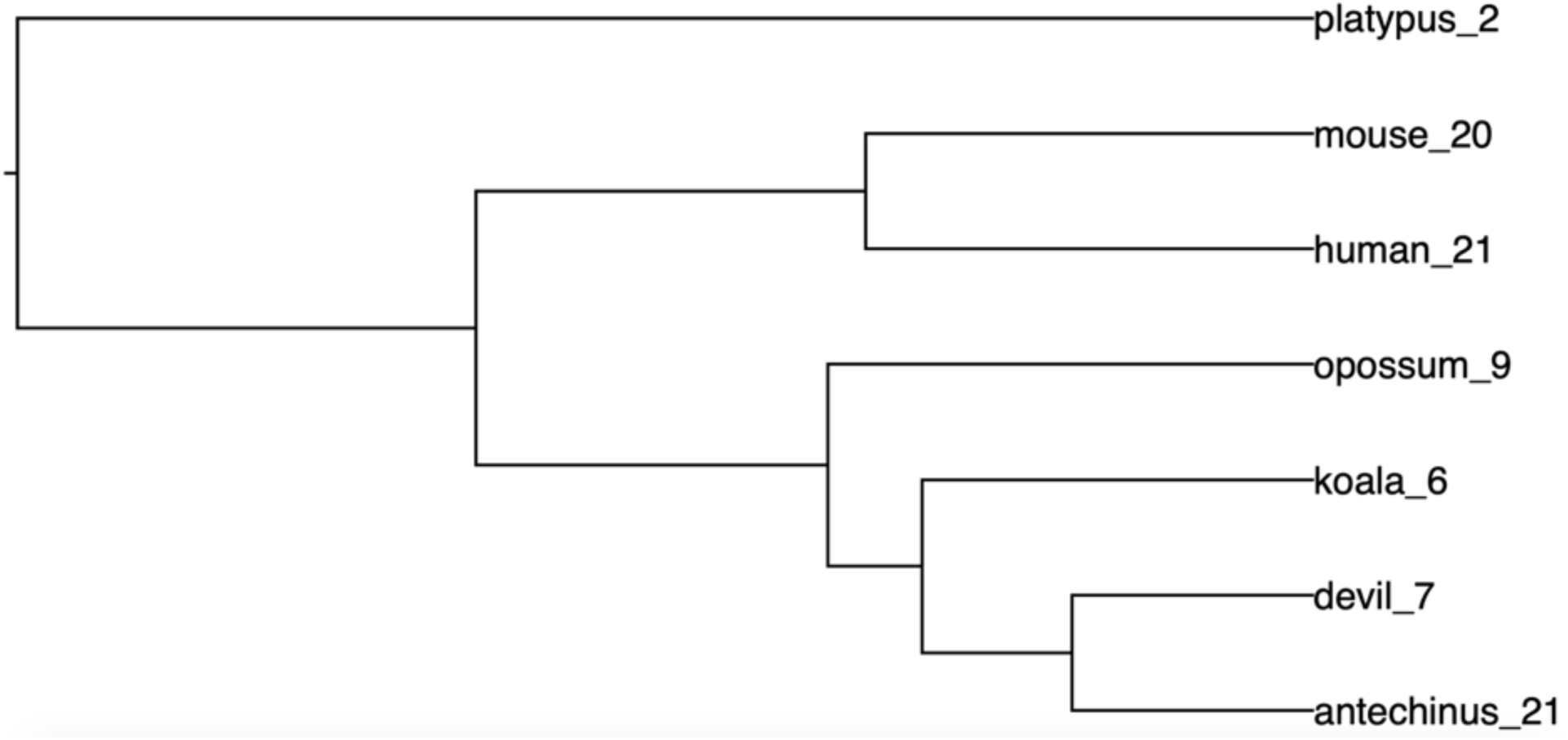
Gene tree showing numbers of Pcdh-γ genes across 7 species. For more information refer to Orthogroup OG0000022 in Supplementary Table 6.

The antechinus was also found to contain a significant expansion of the NMRK2 gene which appears to be single copy in each of the other species. The NMRK2 gene (Nicotinamide Riboside Kinase 2) is involved in the production of NAD+ (Nicotinamide Adenine Dinucleotide), an essential co-enzyme for various metabolic pathways [83, 84]. The antechinus contains 11 full-length copies of this gene in its genome (Figure 7). Furthermore, genes encoding the subunits of the NADH dehydrogenase enzyme which is responsible for conversion of NADH to NAD+, were among the most highly expressed genes within the antechinus transcriptome across a variety of tissue types (Supplementary Table 7). Declining levels of NAD+ have been associated with aging, suggesting that NAD+ may be a key promoter of longevity [84]. NAD+ has also been associated with Alzheimer’s disease whereby increased levels of the molecule may be a protective factor of the disease [85]. The antechinus collected in the current study were collected just prior to the annual breeding season and were therefore mature adults. However, the observed neuropathologies in antechinus species are found to be most prominent in post-breeding individuals and so the data presented here will provide a useful comparison for future studies that explore the development of these pathologies and associated genetic changes across the breeding season. Further investigations into the unique expansion of NMRK2 genes in the antechinus may provide crucial insights into aging and age-related dementias in humans.

**Figure 7.**
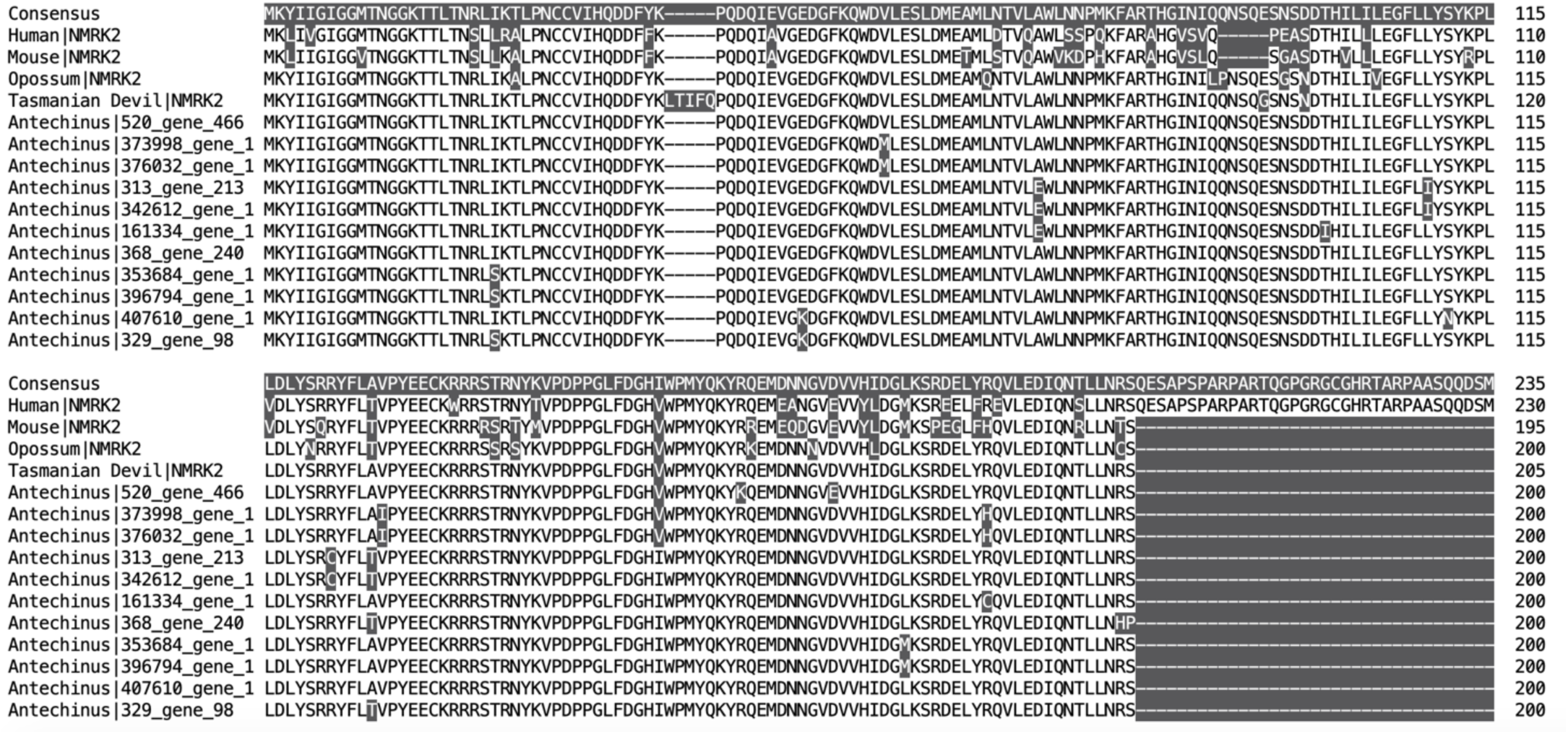
Protein sequence alignment showing expansion of NMRK2 genes in the antechinus. Single copy genes in the human, mouse, gray short-tailed opossum and Tasmanian devil are shown for comparison.

### Alzheimer’s Genes Analysis

To investigate further the potential of antechinus being a disease model for AD [3, 9], we analysed expression and identified variation in genes that have previously been associated with AD. Of the 40 target Alzheimer’s-associated genes, 39 were annotated in the male antechinus reference genome and all 40 were found to be expressed in the global transcriptome (Table 3). The CD2AP gene was not annotated by Fgenesh++ so was not included in downstream analysis. 33 of the annotated antechinus proteins were homologous to human proteins with similarity > 30% [86] (Table 3). Of the seven antechinus gene annotations that showed poor similarity to humans, three (SORL1, CLNK and SLC24A4) were found to have homologous protein-coding transcripts in the global transcriptome suggesting the genome annotations were poor for these genes (likely due to gaps in the reference genome) (Table 3). The remaining four genes (CD33, ZCWPW1, ABCA7 and CR1) did not have homologous genome annotations or transcripts in the antechinus (large gaps were displayed in all sequences compared to the human genes) and were therefore excluded from downstream analysis.

Six of the target genes, including APP, PICALM, KAT8, APOE, INPP5D and MAPT were within the top 90% most highly expressed genes of the global transcriptome and were all found to be expressed in the brain (Supplementary Table 7). Of these genes, APP (amyloid precursor protein) showed the highest level of expression in antechinus brain tissue (Supplementary Table 7). APP is the precursor for the amyloid beta (Aβ) proteins that form amyloid plaques in the brain and is predicted to contribute to early-onset AD in humans [87]. The MAPT gene was also most highly expressed in antechinus brain tissue (Supplementary Table 7) and is responsible for the creation of tau proteins which form the neurofibrillary tangles associated with AD [88]. APOE (apolipoprotein E) is the most common risk-factor gene associated with late-onset AD [89] and was highly expressed across a range of antechinus tissues including the brain (Supplementary Table 7). PICALM is another common gene which has been associated with an increased risk of developing late-onset AD [90]. PICALM is predicted to help flush Aβ proteins out of the brain and so increased expression of the PICALM gene in the brain is predicted to reduce AD risk [91]. This gene was found to be quite lowly expressed in antechinus brain tissue when compared with other tissues such as the spleen or in the blood (Supplementary Table 7) suggesting that it may be contributing to the development of Aβ plaques observed in the antechinus. Finally, KAT8 and INPP5D have been linked to AD through genome-wide association studies [92, 93] and may also be candidates for downstream research. Our finding of expression of some of the most common AD-associated genes in the antechinus brain confirm the potential for this species to be utilized as an AD disease model.

A large variety of genetic variants have been associated with AD in humans, primarily due to their impact on gene expression [92, 94-98]. We utilised the annotated genome-wide SNV data to determine whether antechinus also exhibit variation at Alzheimer’s-associated genes. A total of 16,761 high-quality SNVs (which passed all of the 10x Genomics filters) were associated with the 40 target genes with majority of these being intronic (Figure 8). A total of 81 phased nonsynonymous SNVs were identified across 20 of the target genes, of which 24 were genotyped in both the male and female (Figure 8c, Supplementary Table 8). While the phenotypic effects of these putatively functional variants are currently unknown, mutations in these genes are commonly associated with AD neuropathologies in humans [92, 94-98] and may also be associated with the age-related development of neuropathologies observed in mature antechinus brains [3].

**Figure 8.**
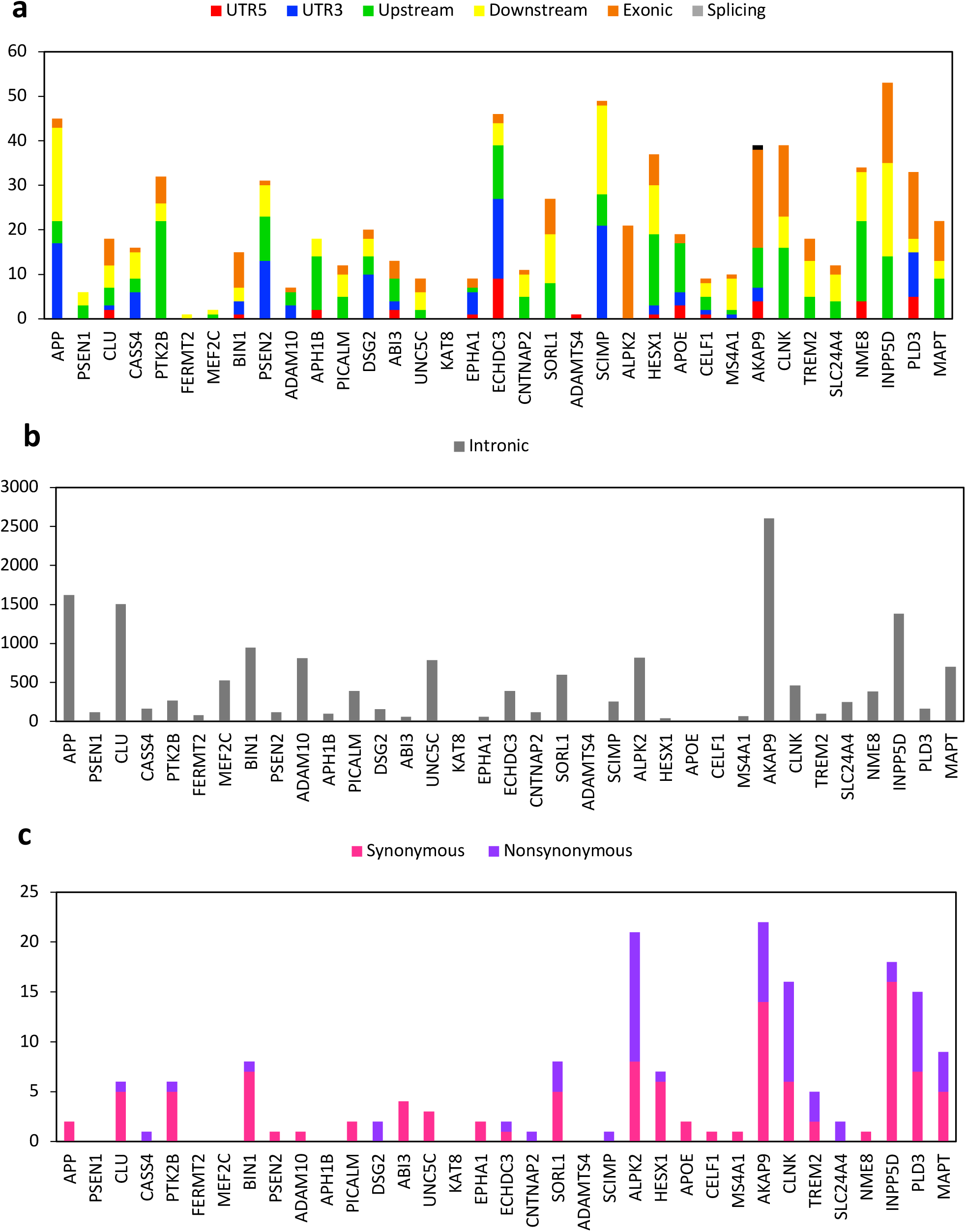
Number of each type of SNV associated with the target Alzheimers-related genes in the antechinus. **a)** Numbers of SNVs present in the 5’ UTR, 3’ UTR, 1kb upstream region, 1kb downstream region, exons, and splice sites of each gene. **b)** Numbers of intronic SNVs present in each gene. **c)** Number of synonymous and nonsynonymous SNVs present in each gene.

## Conclusions and Implications

Here we present the first annotated reference genome within the antechinus genus for a common species, the brown antechinus. The reference genome assembly exhibits completeness comparable to the two current most high-quality marsupial assemblies available (Tasmanian devil and koala), and contains the largest amount of Y-chromosome sequence identified in a marsupial species. Characterisation and annotation of phased, genome-wide variants (including large structural variants) demonstrates considerable diversity within the brown antechinus and provides a resource of gene regions that may have functional implications both in this antechinus and closely related species. Gene ontology analysis of the annotated antechinus proteins identified genes involved in a wide range of biological processes such as immunity, reproduction and stress demonstrating the value of this reference genome in supporting future work investigating the genetic interplay of such processes in this semelparous species. A comparative analysis revealed a number of fast-evolving gene families in the antechinus, most notably within the protocadherin gamma family and NMRK2 gene which have previously been associated with aging and/or aging-related dementias. Target gene analysis revealed high levels of expression of some of the most common genes associated with Alzheimer’s disease in the brain, as well as a number of associated variants that may be involved in the Alzheimer’s-like neuropathological changes that occur in antechinus species. Future research will be able to use the antechinus genome as a springboard to study age-related neurodegeneration, as well as a model for extreme life history trade-offs like semelparity.

## Availability of Supporting Data and Materials

All raw sequencing reads and the draft reference genome have been submitted to NCBI under the BioProject accession PRJNA664282.

## Additional Files

**Supplementary Table 1**: Summary of version 3 and version 4 mammalian BUSCO scores for the antechinus genome and transcriptome assemblies.

**Supplementary Table 2**: Antechinus chromosome assignment comparison based on the AD-ratio approach and mapping to the Tasmanian devil genome.

**Supplementary Table 3**: BLASTP results of annotated antechinus proteins against the Swiss-Prot database using an evalue cut-off of 1e-5 and only reporting the top hit for each gene. **Supplementary Table 4**: Summary of large high-quality structural variants identified in the male and female antechinus and associated genes. Refer to the structural variant VCF files for additional information.

**Supplementary Table 5:** Gene ontology (GO) annotation of annotated antechinus proteins using the generic GO Slim subset. Refer to Supplementary Table 3 to retrieve the antechinus gene annotation IDs for the respective Swiss-Prot gene matches.

**Supplementary Table 6**: Summary of fast evolving gene families including the number of genes in each family across species, the change in the number of genes compared with the predicted ancestral node, the probability of this change, and the respective gene IDs. Numbers represent tree nodes based on the following ultrametric rooted species tree in newick format: (platypus<11>:0.292961,((human<7>:0.101184,mouse<6>:0.101184)<9>:0.0881084,(opossum<5>:0.109848,((antechinus<1>:0.0546286,devil<0>:0.0546286)

**Supplementary Table 7**. Summary of the top 90% most highly expressed genes in the antechinus global transcriptome. For instances where multiple transcripts matched to the same Swiss-Prot gene, the most highly expressed transcript was selected.

**Supplementary Table 8**: Summary of the nonsynonymous SNVs identified in the target Alzheimer’s related genes. Refer to the annotated merged vcf file for additional information.

## Abbreviations

AD: Alzheimer’s disease
RNA: ribonucleic acid
miRNA: microRNA
DNA: deoxyribonucleic acid
SNV: single nucleotide variant
HMW: high molecular weight
bp: base pairs
kb: kilobase pairs
Mb: megabase pairs
Gb: gigabase pairs
PE: paired-end
BUSCO: Benchmarking Universal Single-Copy Orthologs
AD-ratio: average depth ratio
BLAST: Basic Local Alignment Search Tool
NCBI: National Center for Biotechnology Information
BED: Browser Extensible Data
VCF: Variant Call Format
CDS: coding domain sequence
ANNOVAR: Annotate Variation
CAFE: computational analysis of gene family evolution
CNV: copy number variant
SV: structural variant
SNP: single nucleotide polymorphism

## Declarations

### Ethics Statement

All samples were collected in accordance with the *Animal Research Act 1985, Animal Research Regulation 2010*, the *Australian code for the care and use of animals for scientific purposes 8th edition 2013* (the Code) and the *Biodiversity Conservation Act 2016*. University of Sydney Animal Ethics Committee number: 2018/1438 and NSW Scientific License number SL101204.

## Authors’ Contributions

P.B., K.B. and C.H. conceived and designed the project. K.B. and C.H. provided funding. P.B., C.H. and R.S.P.J. collected the samples, P.B prepared the samples, and P.B. and S.T. analysed the data. P.B drafted the manuscript. S.T, C.H, R.S.P.J. and K.B modified the manuscript. All authors read and approved the final version of the manuscript.

## Acknowledgements

We thank Emma Peel for her assistance with DNA and RNA extractions and advice on sample collection. We thank Peter Banks and Mathew Crowther for their guidance on trapping procedures and the Desert Ecology Research Group at the University of Sydney for providing us with the necessary trapping equipment. This research was supported by the Sydney Informatics Hub and the Australian BioCommons which is enabled by NCRIS. Compute resources were provided through a University of Sydney partnership with RONIN and AWS (Amazon Web Services) with the support of Intel.

